# Deafness rapidly reorganizes functional brain networks in adult mice

**DOI:** 10.64898/2026.04.30.721913

**Authors:** Hyun-Ji Shim, Won Beom Jung, Geun Ho Im, Sangyeol Lee, Gunsoo Kim, Seong-Gi Kim

## Abstract

Sensory loss triggers crossmodal reorganization across sensory modalities, and accumulating evidence indicates that this adaptive capacity persists into adulthood. However, the global organizing principles of such plasticity remain poorly understood, as conventional animal model approaches do not permit longitudinal, whole-brain measurements. Here, we use ultra-high-field (15.2T) BOLD fMRI to map deafening-induced functional reorganization across the entire brain in young adult mice. Within one week of deafening, the auditory cortex is recruited by somatosensory and visual inputs, while stimulus-evoked responses are potentiated in the spared sensory pathways. Reorganization extends beyond sensory cortices to higher-order association areas, including anterior cingulate, retrosplenial, and posterior parietal cortices. Resting-state fMRI further reveals strengthened coupling both within sensory systems and between sensory systems and a default mode-like network. These findings demonstrate that adult-onset deafness rapidly reorganizes functional brain networks and further implicate the default mode-like network as a potential mediator of crossmodal integration.

## Introduction

Sensory loss triggers functional reorganization across sensory pathways^1,2^. In congenitally or early-deaf individuals, visual and tactile stimulation recruit deprived auditory cortical areas^3–5^. This crossmodal plasticity is thought to underlie behavioral enhancements, including superior visual motion processing^6–10^ and increased tactile sensitivity^11^.

Although plasticity is generally considered more constrained in adulthood, accumulating evidence indicates that hearing loss acquired later in life can still drive substantial crossmodal reorganization. Humans with late-onset hearing loss exhibit visual activation within auditory cortical regions, as shown by visually evoked potentials^12–16^ and fMRI responses^17^. Similarly, electrophysiological recordings in animal models have shown that adult-onset deafness induces somatosensory responses in the auditory cortex^18,19^ and increases the proportion of visually responsive neurons in secondary auditory areas^20^. Beyond such crossmodal recruitment of the deprived auditory cortex, adult-onset deafness potentiates thalamocortical synaptic connections in the visual cortex^21,22^ and visually evoked potentials^23^, suggesting that spared sensory pathways are reshaped as well^24^.

Despite this growing recognition, the full scope and global organizing principles of adult-onset crossmodal plasticity remain poorly understood. Prior work, largely using visual deprivation models, has emphasized local connectivity changes mediated by homeostatic and synaptic plasticity as the primary drivers of deprivation-induced reorganization^24^. However, sensory cortices are extensively interconnected with subcortical and higher-order cortical networks^25,26^, and sensory processing is shaped by large-scale intrinsic dynamics, including interactions with the default mode network (DMN)^27,28^. Sensory loss may therefore alter network-level interactions^29,30^, driving reorganization well beyond local synaptic adjustments. Yet, few animal studies have examined functional plasticity at the whole-brain level, leaving it unclear whether adult-onset deafness reorganizes large-scale functional networks.

Functional magnetic resonance imaging (fMRI) provides a powerful tool for probing brain-wide network dynamics and plasticity. By measuring blood-oxygenation level dependent (BOLD) signals associated with neural activity^31–33^, fMRI enables repeated, non-invasive assessment of stimulus-evoked responses across subcortical structures, sensory and higher-order cortical regions simultaneously, overcoming limitations of region-specific analyses restricted to individual sensory areas. Furthermore, resting-state fMRI allows characterization of task-independent functional networks^27,34,35^, enabling investigation of how large-scale intrinsic network dynamics are reshaped following sensory loss.

Here we performed longitudinal high-field BOLD fMRI in genetically engineered mice before and after deafening to map brain-wide changes in both stimulus-evoked and intrinsic network activity. We find that adult-onset deafness potentiates spared sensory pathways while driving crossmodal recruitment of auditory cortex and higher-order cortical regions. Resting-state fMRI further reveal enhanced functional connectivity within and between sensory, motor, and DMN-like networks. Together, these results demonstrate that adult-onset deafness rapidly reorganizes neural circuits at multiple scales, reconfiguring large-scale network connectivity alongside local plasticity within sensory cortex.

## Results

### Probing deafening-induced plasticity with longitudinal BOLD fMRI in adult mice

To investigate how adult-onset deafness reshapes brain-wide functional organization, we performed longitudinal ultra-high-field (15.2 T) BOLD fMRI in young adult mice (Fig. 1). Deafness was induced in Pou4f3^DTR/+^ (DTR) mice by diphtheria toxin (DT) injection, which ablates cochlear hair cells (Fig 1Bb)^36,37^. Deafness was confirmed one week after DT treatment by the absence of acoustic startle responses (ASR) in all mice prior to imaging (Fig. 1Ba). BOLD fMRI was conducted in the same animals before and one week after DT treatment to examine deafening-induced changes in evoked sensory responses as well as intrinsic functional connectivity across the brain (Fig. 1C).

**Figure 1.**
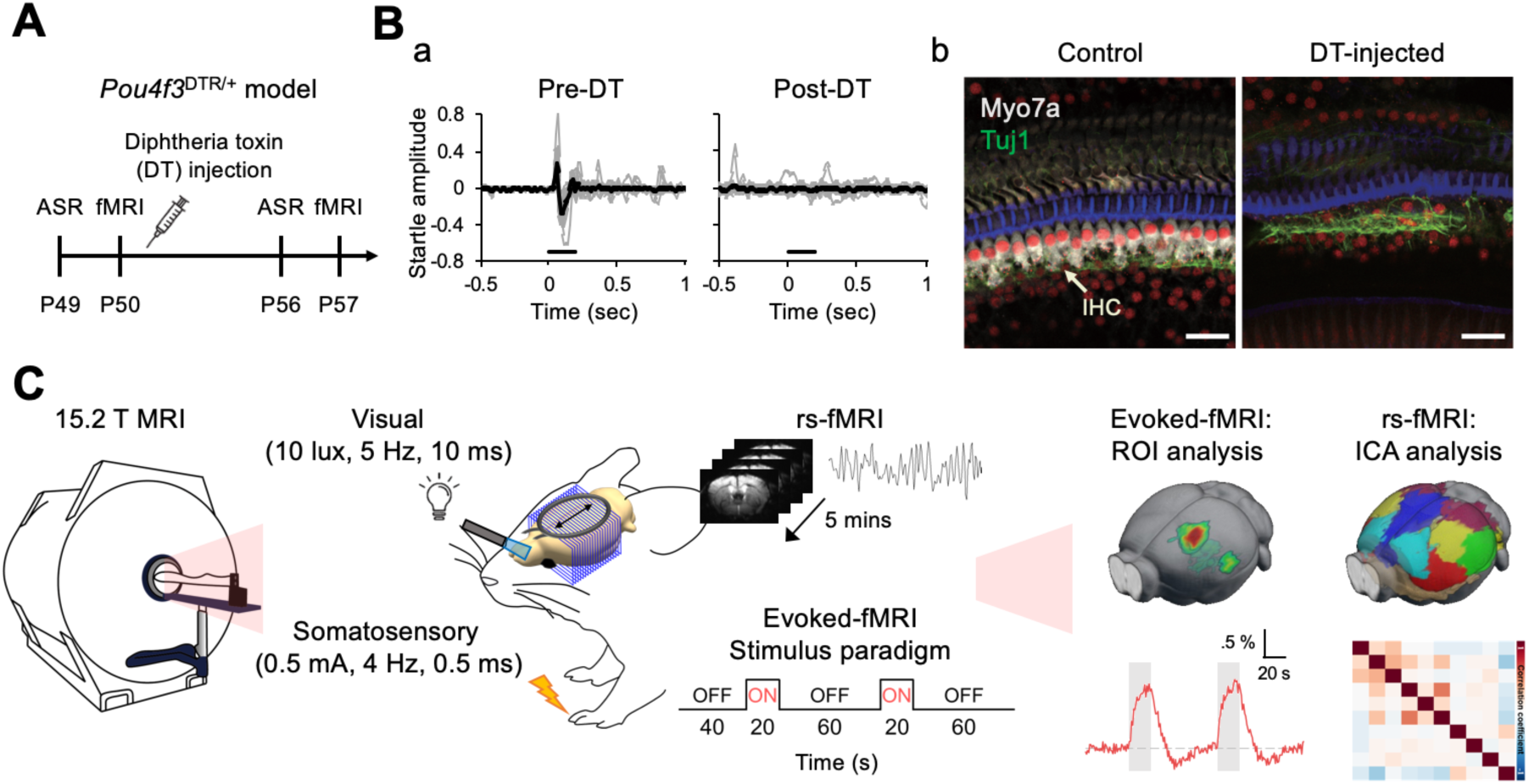
Experimental design for deafening and longitudinal BOLD fMRI. (A) Experimental timeline showing diphtheria (DT) injection and subsequent acoustic startle response (ASR) testing and fMRI sessions. (B) Validation of deafening in Pou4f3^DTR/+^ (DTR) mice. a. Example ASR traces from an individual mouse before and after DT treatment, demonstrating loss of startle responses following deafening. b. Whole mount cochlear histology from an untreated DTR mouse and a DT-injected DTR mouse. Hair cells are labeled in gray (Myosin VIIa), auditory nerve fibers in green (Tuj1), and hair cell cilia and scaffold proteins in blue (phalloidin), showing near-complete hair cell ablation after DT treatment. Scale bar, 20 µm. (C) Overview of the ultra-high field (15.2 T) BOLD fMRI experimental setup. Somatosensory (forepaw electrical stimulation) and visual (light delivered via optic fibers) stimulation paradigms for evoked fMRI are shown, along with the resting-state fMRI (rs-fMRI) acquisition protocol. Analysis pipelines for evoked fMRI (ROI-based analysis) and rs-fMRI (independent component analysis) are shown schematically.

To examine how adult-onset deafening influences spared sensory pathways, we measured BOLD responses evoked by somatosensory and visual stimulation. Somatosensory responses were elicited by unilateral electrical forepaw stimulation, interleaved with bilateral visual stimulation delivered via optical fibers.

Representative activation maps and response time courses from individual mice show that, prior to deafening, both WT and DTR mice exhibited reliable somatosensory-evoked BOLD responses in the contralateral primary somatosensory cortex (S1) (Fig. 2A, black traces), consistent with previous reports^38–40^. Visual stimulation similarly evoked robust BOLD responses in primary visual cortex (V1) before deafening (Fig. 2B, black traces).

**Figure 2.**
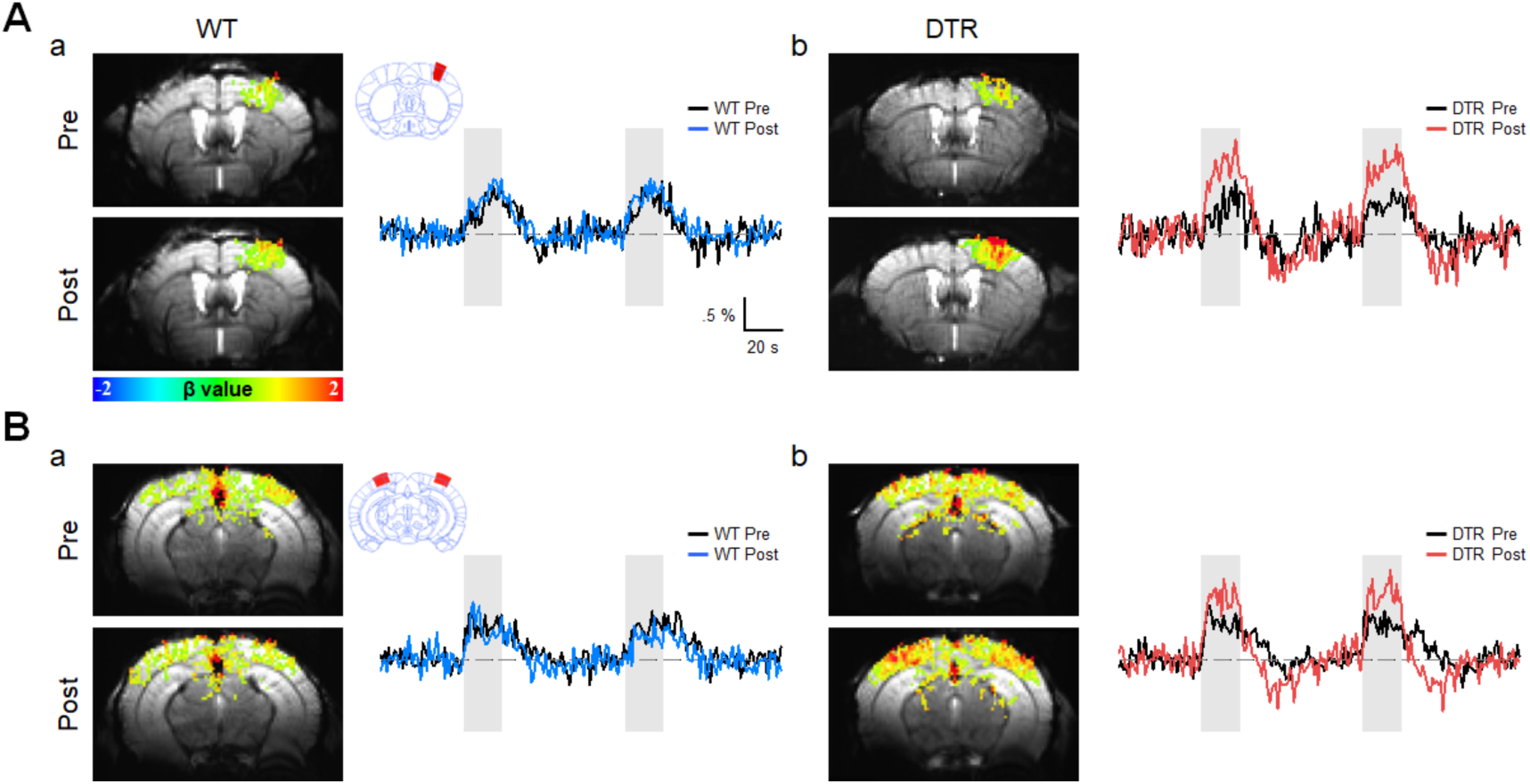
Somatosensory- and visual-evoked BOLD fMRI responses in individual mice. (A) Somatosensory BOLD responses evoked by left forepaw stimulation in the right primary somatosensory cortex (S1) of a WT (a) and a DTR (b) mouse. Representative BOLD activation maps (β values) are shown before (Pre) and after (Post) DT treatment. Corresponding BOLD time courses averaged within the S1 ROI are shown to the right. (B) Visual BOLD responses evoked by bilateral stimulation in the primary visual cortex (V1) of the same WT (a) and DTR (b) mice. Representative activation maps and corresponding ROI-averaged BOLD time courses are shown before and after DT treatment. Black traces indicate responses before DT; blue (WT) and red (DTR) traces indicate responses after DT treatment. Grey shaded areas denote sensory stimulation periods.

In the example DTR mouse, somatosensory responses were enhanced one week after DT treatment (Fig. 2A, red trace), whereas the example WT mouse showed stable response amplitudes across sessions (Fig. 2A, blue trace). Visual-evoked responses were also increased in the DTR mouse (Fig. 2B, red trace) but remained unchanged in the WT mouse (Fig. 2B, blue trace). These representative examples demonstrate that deafening can induce robust enhancement of BOLD responses in spared primary sensory cortices.

### Enhanced somatosensory responses and cross-modal cortical recruitment after deafening

To investigate the consistency of sensory enhancements across animals, voxel-wise activation maps from individual mice were registered to a reference atlas (see Methods). Fifteen regions of interest (ROIs) were then defined according to the atlas for quantitative analysis of response amplitudes in each animal (Fig. 3Aa). Average activation maps show that prior to deafening, both WT and DTR mice exhibited robust activation along canonical somatosensory pathways, including primary and secondary somatosensory cortex (S1, S2), motor cortex (M1), and the ventral posterior thalamic nucleus (VP) (Fig. 3A).

**Figure 3.**
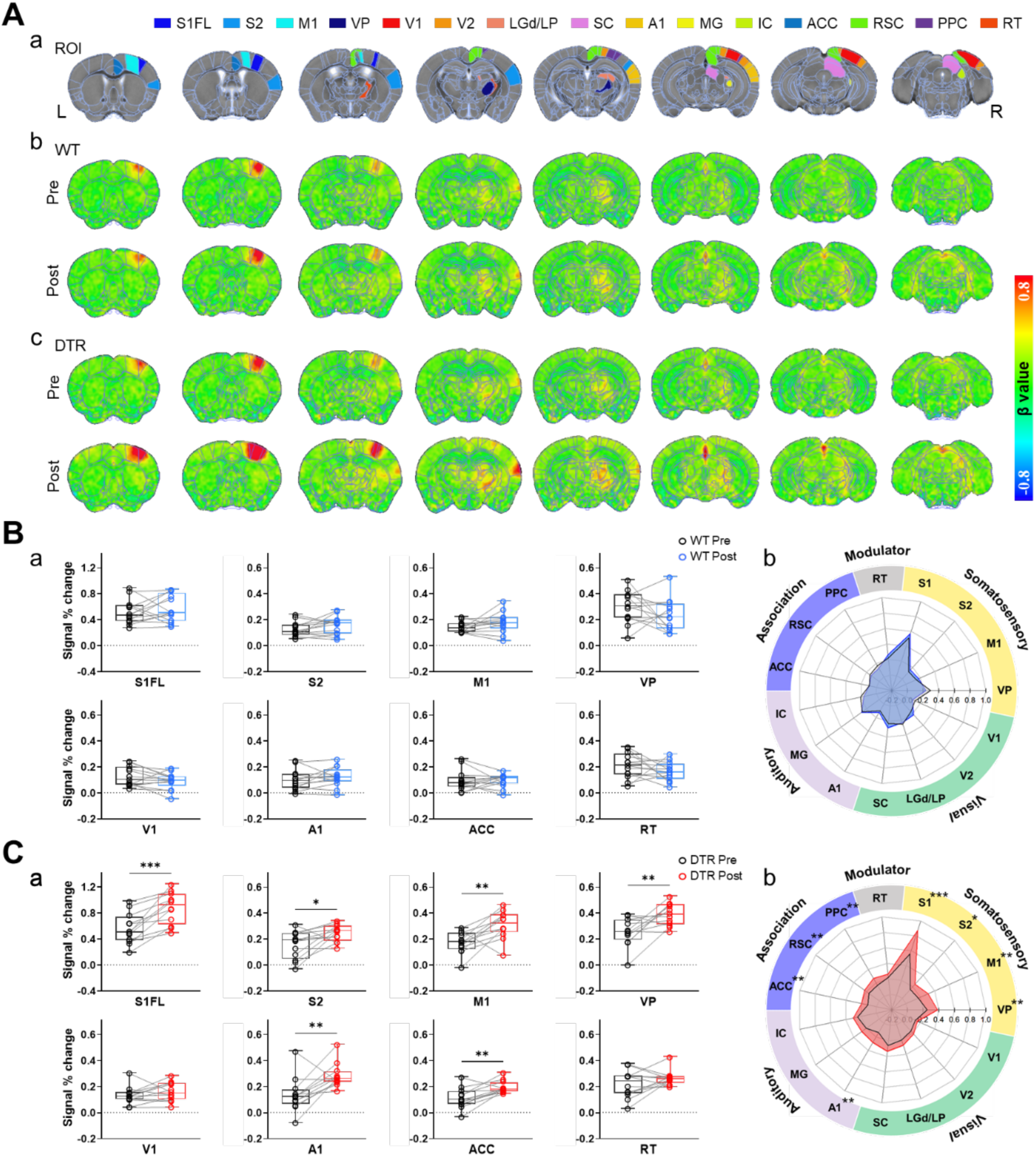
Cross-modal enhancement of somatosensory-evoked BOLD responses across brain areas after deafening. (A) Group-averaged activation maps during forepaw stimulation. a. Anatomical reference images overlaid with Allen Brain Atlas-based regional parcellation are shown across eight coronal slices (anterior to posterior). Fifteen regions of interest (ROIs), spanning somatosensory, visual, auditory, association, and reticular thalamus, are displayed in distinct colors. Group-averaged BOLD activation maps (β values) are shown for wild-type (WT; b) and Pou4f3^DTR/+^ (DTR; c) mice before (Pre) and one week after (Post) diphtheria toxin (DT) injection. (B) Quantification of somatosensory-evoked BOLD responses in WT mice. a. ROI-based response amplitudes measured before and after DT injection for selected regions (S1FL, S2, M1, VP, V1, A1, ACC, RT). Each data point represents an individual animal, with lines connecting pre- and post- measurements. b. Circular summary plots illustrate response amplitudes across all 15 ROIs, grouped by functional system. WT mice show stable responses across all regions (n = 13 mice, p > 0.05). S1FL: forelimb region of S1; VP: ventral posterior complex of the thalamus; ACC: anterior cingulate cortex; RSC: retrosplenial cortex; PPC: posterior parietal cortex. MG: medial geniculate body; IC: inferior colliculus. LGd: dorsal part of the lateral geniculate complex; LP: lateral posterior nucleus of the thalamus; SC: superior colliculus; RT: reticular thalamus. (C) Quantification of somatosensory-evoked BOLD responses in DTR mice. a. ROI-based analyses reveal significant post-deafening increases in the somatosensory cortex (S1FL, S2), motor cortex (M1), thalamus (VP), auditory cortex (A1), and ACC, whereas the visual cortex (V1) and reticular thalamus (RT) show no significant change. b. Circular summary plots show enhanced responses across somatosensory and association cortices, as well as recruitment of auditory cortex following deafening (n = 12 mice). * p < 0.05, ** p < 0.01, *** p < 0.001, paired t-test. Plots without asterisks indicate no significant differences (p > 0.05).

One week after DT treatment, WT mice showed stable somatosensory responses, with no significant changes in either cortical or subcortical regions (Fig. 3B; Supplementary Fig. 1A and Table 1). In contrast, DTR mice displayed markedly enhanced somatosensory-evoked BOLD responses across the same cortical and thalamic regions (Fig. 3A). Quantitative within-animal, ROI-based comparisons confirmed significant post-deafening increases in response amplitudes in S1, S2, M1, and VP relative to pre-deafening baseline levels (Fig. 3C; Supplementary Fig. 1B and Table 2).

Deafening also enhanced somatosensory responses in the primary auditory cortex (A1) and in higher-order association cortical areas, including anterior cingulate cortex (ACC), retrosplenial cortex (RSC), and posterior parietal cortex (PPC) – regions that exhibited minimal somatosensory responses prior to deafening (Fig. 3C; Supplementary Table 2). This rapid recruitment of auditory cortex by somatosensory input represents a hallmark of cross-modal plasticity, where deprived sensory regions become driven by spared modalities. The post-deafening enhancement observed in association cortices further suggests that deprivation-induced plasticity extends beyond primary sensory pathways to reorganize distributed cortical networks.

These findings indicate that adult-onset deafness rapidly enhances intramodal somatosensory processing across both cortical and subcortical pathways while concurrently recruiting deprived auditory cortex and higher-order cortical regions.

### Enhanced visual responses and cross-modal cortical recruitment after deafening

To determine whether deafening broadly affects other spared sensory modalities, we also examined visual-evoked BOLD responses in the same animals. Prior to DT treatment, visual stimulation reliably activated the primary and secondary visual cortices (V1, V2), as well as visual thalamic nuclei (LGd and LP) and the superior colliculus (SC) in both WT and DTR mice (Fig. 4A), consistent with previous findings^41^. One week after DT treatment, WT mice exhibited stable visual responses with no significant changes in either cortical or subcortical regions (Fig. 4B; Supplementary Fig. 2A and Table 3).

**Figure 4.**
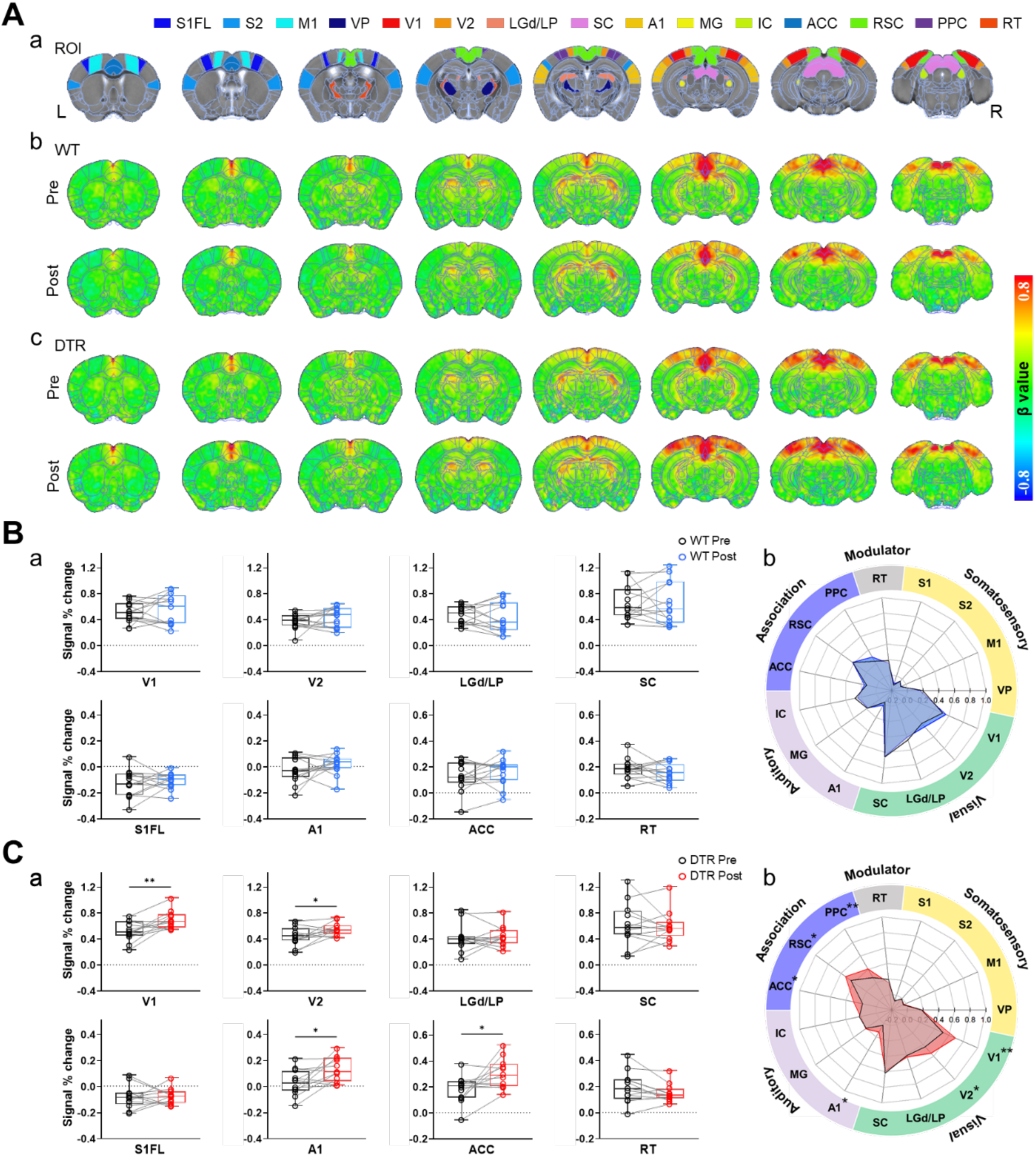
Cross-modal enhancement of visual-evoked BOLD responses across brain areas after deafening. (A) Group-averaged activation maps during visual stimulation. a. Anatomical reference images overlaid with Allen Brain Atlas-based regional parcellation are shown across eight coronal slices (anterior to posterior). Fifteen regions of interest (ROIs) spanning visual, somatosensory, auditory, association, and control regions are displayed in distinct colors. Group-averaged BOLD activation maps (β values) are shown for WT (b) and DTR (c) mice before (Pre) and one week after (Post) DT injection. (B) Quantification of visual-evoked BOLD responses in WT mice. a. ROI-based response amplitudes measured before and after DT injection for selected regions (V1, V2, LGd/LP, SC, S1FL, A1, ACC, RT). Each data point represents an individual animal, with lines connecting pre- and post-DT treatment measurements. b. circular summary plots illustrate response amplitudes across all 15 ROIs grouped by functional system. WT mice show stable visual responses across all regions (n = 13 mice). (C) Quantification of visual-evoked BOLD responses in DTR mice. a. ROI-based analyses reveal significant post-deafening increases in primary and secondary visual cortex (V1, V2), auditory cortex (A1), and ACC, whereas visual subcortical nuclei (LGd/LP, SC), somatosensory cortex (S1FL), and retinular thalamus (RT) show no significant change. b. Circular summary plots show enhanced responses across visual and association cortices and crossmodal recruitment of auditory cortex following deafening (n = 12 mice). * p < 0.05, ** p < 0.01, *** p < 0.001, paired t-test. Plots without asterisks indicate no significant differences (p > 0.05).

Following deafening, visual stimulation evoked significantly enhanced responses in V1 and V2 of DTR mice, as confirmed by ROI-based quantification (Fig. 4C; Supplementary Fig. 2B and Table 4). In contrast to somatosensory stimulation, enhancement of visual-evoked responses was restricted to cortical regions, with no significant changes detected in visual thalamic nuclei or the superior colliculus. As observed with somatosensory stimulation, visual stimulation after deafening also recruited the auditory cortex and association cortical regions, including the ACC, RSC, and PPC (Fig. 4C). Interestingly, deafening enhanced activity in prefrontal and association cortices during both visual and somatosensory stimulation, indicating that loss of auditory input reorganizes higher-order networks, including the default mode network (DMN)-like regions such as ACC and RSC.

Together, these results demonstrate that adult-onset deafness induces modality-specific enhancement in spared sensory pathways while enabling cross-modal recruitment of deprived auditory cortex and engaging higher-order association networks.

### Enhanced sensory responses are not explained by MR scanner noise or cross-modal cortical suppression

We next asked whether the observed enhancement in spared sensory modalities could be attributed to factors other than cross-modal plasticity. Two alternative explanations were considered: 1) the effects of high-intensity acoustic MR scanner noise on sensory-evoked responses, and 2) release from cross-modal cortico-cortical suppression.

During BOLD signal acquisition, high intensity acoustic noise is generated by the rapid switching of electric currents in the gradient coils. Previous studies have reported that such scanner noise can modulate BOLD responses in humans and rodents^42,43^. If acoustic noise suppresses somatosensory-evoked responses in hearing mice, elimination of this suppression after deafening could potentially account for the enhanced sensory responses observed in DTR mice.

To test this possibility, we performed optical intrinsic signal (OIS) imaging, which closely parallels BOLD fMRI signals^44^, while presenting playback of recorded MR scanner noise. OIS imaging revealed robust somatosensory activation in S1 with or without noise playback, with comparable spatial extent, response amplitude, and temporal dynamics (Fig. 5A-C). Within-animal comparisons showed no significant difference in response amplitude between quiet and noise conditions (Fig. 5D; n = 5 mice, p = 0.21, paired t-test). These results indicate that MR scanner noise does not suppress somatosensory responses in hearing mice and therefore cannot explain the post-deafening enhancement.

**Figure 5.**
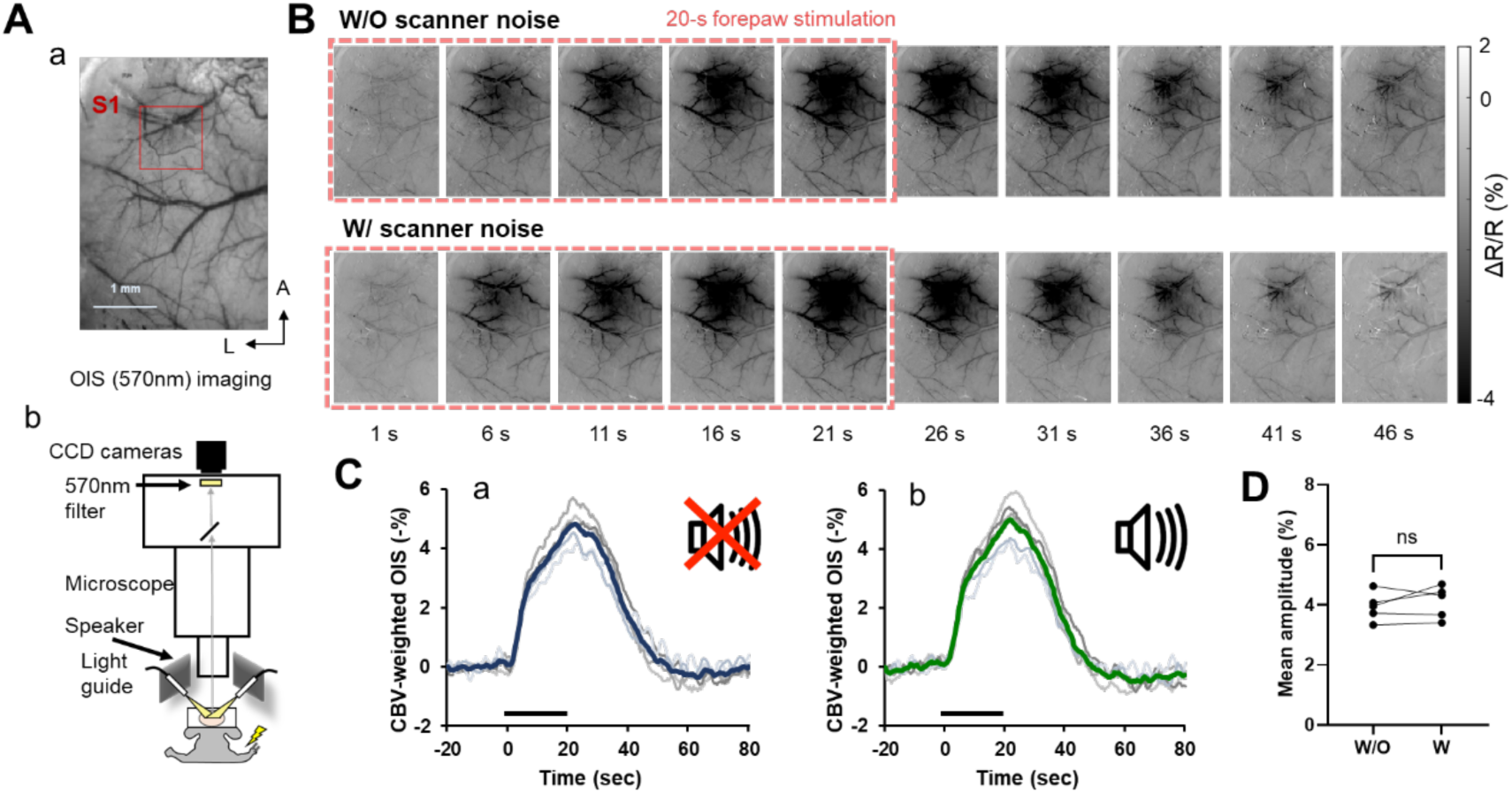
MR scanner acoustic noise does not suppress somatosensory responses. (A) Optical intrinsic signal (OIS) imaging setup and region of interest (ROI) definition. a. Thinned-skull image of the left hemisphere acquired at 570 nm, with a square ROI placed over primary somatosensory cortex (S1FL). b. Schematic of the OIS experimental setup showing illumination of the cortical surface, CCD-based reflectance imaging, and speaker delivering recorded MR scanner noise. (B) CBV-weighted OIS activation maps during forepaw stimulation acquired without (top) and with (bottom) scanner noise playback. Images show changes in reflectance (ΔR/R, %) across time, spanning stimulation and recovery periods. Darkening indicates increased total hemoglobin concentration. (C) Time courses of somatosensory-evoked OIS responses extracted from the S1 ROI in the absence (a) and presence (b) of scanner noise playback. Thin lines represent individual animals, and thick lines indicate the group mean. (D) Quantification of peak somatosensory response amplitudes with and without scanner noise. Each data point represents an individual animal, with lines connecting paired measurements. No significant difference was observed between conditions (noise-off: 3.94 ± 0.21% vs. noise-on: 4.10 ± 0.24%; n = 5 mice, p = 0.21, paired t-test). ns, not significant.

Cross-modal suppression of cortical sensory responses has been reported and is thought to be mediated, in part, by direct cortico-cortical projections between sensory regions^45–47^. If such cross-modal suppression exists under baseline conditions, elimination of ongoing inhibitory influence from the auditory cortex after deafening could lead to enhanced responses in spared modalities.

We therefore tested whether acute suppression of auditory cortex is sufficient to reproduce the sensory enhancements observed after deafening. In hearing VGAT-ChR2 mice, bilateral optogenetic stimulation of A1 effectively suppressed baseline cortical activity, as confirmed by negative BOLD responses at the stimulation site (Fig. 6A, B). Somatosensory-evoked BOLD responses were then measured during forepaw stimulation with and without continuous bilateral suppression of A1. Somatosensory activation maps and temporal response profiles were indistinguishable between conditions (Fig. 6B-C). Quantitative analysis revealed no significant differences in response amplitudes across somatosensory, motor, thalamic, visual, or auditory cortical regions (Fig. 6D; Supplementary Table 5).

**Figure 6.**
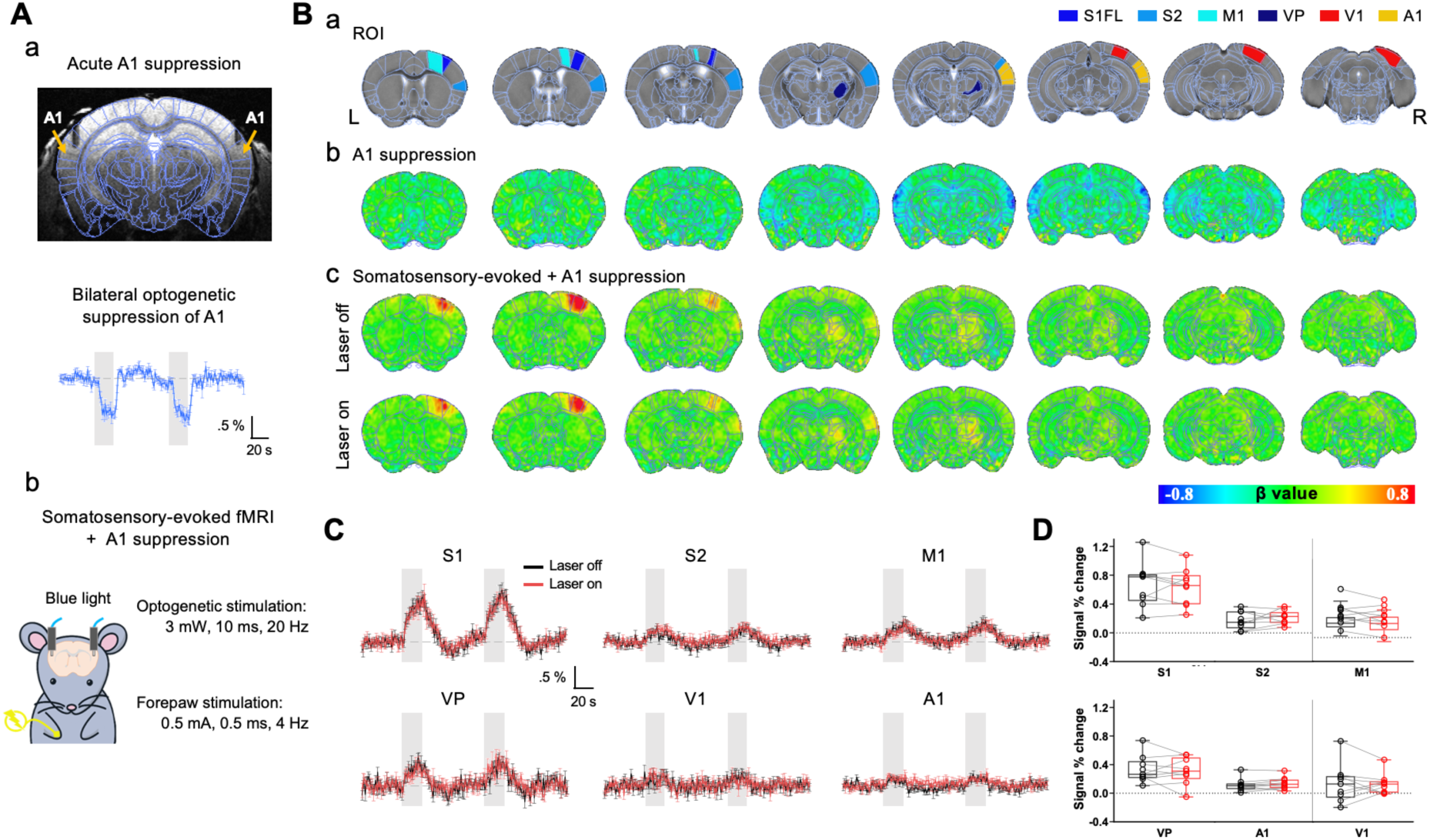
Acute auditory cortex suppression does not reproduce the sensory enhancements observed after deafening. (A) Bilateral optogenetic suppression of A1. a. Anatomical reference image showing bilateral optic fiber placements targeting A1, overlaid with Allen Brain Atlas parcellation. Time courses demonstrate effective optogenetic suppression, indicated by negative BOLD responses during optogenetic stimulation. b. Experimental paradigm for somatosensory-evoked fMRI with continuous bilateral A1 suppression during left forepaw stimulation. Optogenetic stimulation was applied continuously throughout the forepaw stimulation trials. (B) Group-averaged BOLD activation maps. a. Anatomical reference images with selected ROIs (S1FL, S2, M1, VP, V1, A1) indicated. b. Activation maps during optogenetic A1 suppression alone, showing reduced activity confined to auditory cortex. c. Somatosensory-evoked activation maps without (top, laser off) and with (bottom, laser on) concurrent A1 suppression. (C) Time courses of somatosensory-evoked BOLD responses extracted from selected ROIs with (laser on, red traces) and without (laser off, black traces) A1 suppression. Traces overlap across conditions, indicating no detectable modulation by A1 acute suppression. (D) Quantification of somatosensory-evoked response amplitudes across regions. Each data point represents an individual animal, with lines connecting paired measurements. No significant differences were observed between conditions (n = 9 mice, p > 0.05, paired t-test).

Together, these control experiments demonstrate that the enhanced sensory responses following adult-onset deafness cannot be explained by scanner-related acoustic artifacts or acute cortical disinhibition.

### Deafening increases temporal correlations across resting-state functional networks

In addition to enhanced sensory-evoked responses and cross-modal recruitment of auditory cortex by spared sensory inputs following deafening, our results reveal stronger activation of association cortices. Although such changes could arise from local mechanisms, including homeostatic plasticity within individual sensory circuits^24^, the widespread engagement of higher-order cortical regions suggests the involvement of large-scale network reorganization.

Resting-state fMRI can reveal functional brain networks characterized by coordinated spontaneous fluctuations and can be used to assess interactions among these networks^48^. Such interactions may provide insight into how different sensory regions communicate beyond individual sensory modalities. To examine whether deafening alters intrinsic brain functional connectivity, we analyzed resting-state fMRI data acquired before and one week after DT treatment (Fig. 7). Independent component analysis (ICA) applied to resting-state time series signals identified ten large-scale, distributed functional networks corresponding to canonical sensory, motor, default mode, hippocampal, striatal, and olfactory systems (Fig. 7A). These bilaterally organized functional networks were consistent with those reported in previous resting-state fMRI studies in mice^49,50^.

**Figure 7.**
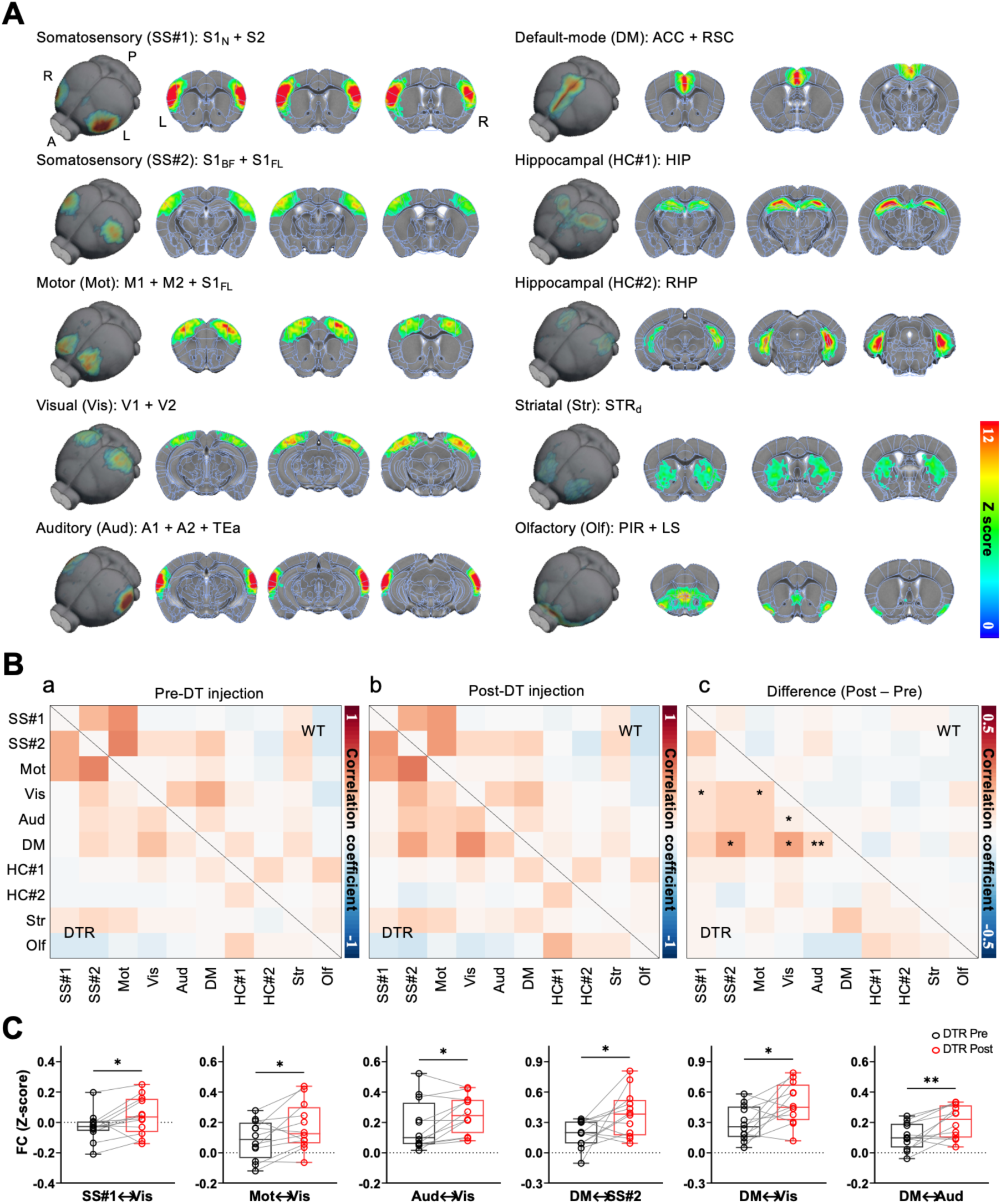
Deafening reorganized resting-state functional connectivity across large-scale networks. (A) Ten resting-sate networks identified by independent component analysis, representing canonical somatosensory (SS#1, SS#2), motor (Mot), visual (Vis), auditory (Aud), default mode (DM), hippocampal (HC#1, HC#2), striatal (Str), and olfactory (Olf) systems. Networks are displayed as thresholded z-score maps and are bilaterally distributed across hemispheres. Three-dimensional renderings and corresponding two-dimensional slices illustrate the spatial extent of each component. (B) Group-averaged functional connectivity matrices showing pairwise correlations between resting-state networks before (Pre) and one week after (Post) DT injection. Left lower triangles correspond to DTR mice, whereas right upper triangles correspond to WT control mice. Matrices are shown for pre-DT (a), post-DT (b), and their difference (c). In difference matrices (c), network pairs with significant changes in correlation are marked with asterisks. Color scale indicates Fisher z-transformed correlation coefficients. (C) Quantification of functional connectivity changes for network pairs showing significant post-deafening increases. Functional coupling increased significantly between somatosensory and visual networks (SS#1-Vis), motor and visual networks (Mot-Vis), auditory and visual networks (Aud-Vis), as well as between default mode network (DM) and somatosensory (SS#2), visual (Vis), and auditory (Aud) networks. Each data point represents an individual animal, with lines connecting pre- and post- deafening measurements. * p < 0.05, paired t-test.

Following DT treatment, WT mice exhibited no systematic changes in functional connectivity (Fig. 7B), indicating that repeated imaging or toxin exposure alone does not significantly influence resting-state connectivity. In contrast, DTR mice showed a broad increase in temporal correlations across resting-state functional networks (Fig. 7B). These increases were most pronounced among sensory networks, between sensory and motor networks, and between sensory networks and a DMN-like network. Specifically, temporal correlations increased significantly between somatosensory and visual networks (SS#1-Vis), motor and visual networks (Mot-Vis), auditory and visual networks (Aud-Vis), as well as between a DMN-like network and multiple sensory networks (DM-SS#2, DM-Vis, DM-Aud) (Fig. 7C; Supplementary Table 6).

These findings indicate that deafening-induced reorganization of functional connectivity extends beyond primary sensory pathways and involves enhanced interactions with higher order cortical regions, including the DMN-like network.

## Discussion

Using longitudinal ultra-high-field BOLD fMRI, we demonstrate that deafening in young adult mice drives robust, brain-wide functional reorganization. Deafening enhances stimulus-evoked responses in spared pathways, recruits the deprived auditory cortex during somatosensory and visual processing, and increases engagement of higher-order association regions. These changes are accompanied by strengthened temporal correlations across large-scale resting-state networks. Together, our findings establish that adult-onset deafness triggers coordinated reorganization across sensory, association, and intrinsic networks, extending well beyond the deprived modality.

### BOLD fMRI mapping of crossmodal plasticity in the mouse brain

BOLD fMRI enables longitudinal, whole brain measurements within the same animal, providing a systems-level view of plasticity that is not readily accessible with spatially restricted electrophysiological or optical approaches. Although BOLD signals reflect hemodynamic rather than neural activity directly, they reliably capture the magnitude and sign of population responses^51–53^, and the potentiation of spared modalities we observe is consistent with strengthened feedforward thalamocortical drive following hearing loss^21,22^.

Potential confounds from MR acoustic noise are unlikely to account for our results^42,43,54,55^. Optical intrinsic signal imaging during scanner noise playback revealed no effect on somatosensory responses, ruling out the possibility that enhanced responses after deafening reflect removal of noise-induced suppression. This may reflect the limited sensitivity of mice to the low-frequency spectral peak (∼1.3 kHz) of echo planar imaging (EPI) noise^56^. These findings support the use of BOLD fMRI as a robust approach for mapping brain-wide reorganization following sensory loss.

### Deafening-induced reorganization of sensory responses across the brain

Deafening induces crossmodal recruitment of the auditory cortex by both somatosensory and visual inputs, consistent with but also extending prior observations in humans and animal models, typically restricted to single modalities^12,16–18,20^. In the adult brain, such recruitment likely arises from strengthening of pre-existing heteromodal inputs from primary and association cortices^24,25^.

Spared modalities are likely potentiated through pathway-specific experience-dependent plasticity rather than a global gain increase. This potentiation is not due to an immediate release from crossmodal suppression, as acute optogenetic suppression of auditory cortex did not reproduce the effect (Fig 6). Somatosensory responses are enhanced across both cortical and subcortical structures, including S1, S2, M1, and somatosensory thalamus (VP), whereas visual enhancement was largely restricted to the cortex. This dissociation demonstrates that adult-onset deafness utilizes different circuit mechanisms to re-weight sensory pathways across the spared modalities. The stronger and more widespread somatosensory effects likely reflect tighter anatomical and functional interactions with auditory circuits^57^.

The presence of thalamic plasticity is consistent with recent evidence for subcortical plasticity following adult-onset sensory loss^26^. Deafening selectively enhances somatosensory thalamic responses without altering visual or auditory thalamic activity. This pathway-specific effect may arise from changes in ascending inputs, partly through modality-specific disinhibition of the primary thalamus^26^, or corticothalamic feedback^58^. In contrast, the absence of crossmodal recruitment in the auditory thalamus suggests that such recruitment primarily occurs in cortical circuits.

### Large-scale network reorganization and DMN engagement

Deafening-induced reorganization extends beyond sensory pathways to large-scale brain networks. Higher-order regions, including ACC, RSC, and PPC, show enhanced responses during sensory stimulation, revealing a systems-level reconfiguration not captured in prior animal studies. This pattern parallels findings in humans with hearing loss and highlights the involvement of association cortices in crossmodal plasticity^16^.

Resting-state fMRI has been widely used to define canonical functional brain networks, including the default-mode network, in both humans and rodents^27,34,35,49,50^. Using this approach, we find that deafening broadly strengthens functional connectivity across large-scale brain networks. Increases were most pronounced within sensory networks, between sensory and motor networks, and between sensory networks and a DMN-like network. While homeostatic models account for synaptic plasticity within individual circuits, they do not explain coordinated changes across distributed networks. The increased functional connectivity between the DMN-like network and sensory networks suggests a higher-order role for association regions, including ACC and RSC, in coordinating activity across modalities following sensory loss. Such global mechanisms likely operate in concert with local synaptic mechanisms to support adaptive reorganization.

### Time course of reorganization and functional implications

The rapid emergence of large-scale changes one week after deafening indicates that adult plasticity operates on short time scales. Prior studies suggest that components of this reorganization arise within days^24,59,60^. Thus, some aspects of the reorganization may occur earlier than observed here.

It remains to be determined whether the observed local and large-scale changes are long-lasting. Crossmodal recruitment and enhancement of spared modalities can persist for months in animal models^18,23^. At the same time, studies in human cochlear implant users indicate that deprived auditory regions remain plastic and responsive to restored input^61^. Therefore, while many aspects of reorganization may be stable, adult sensory circuits probably remain sensitive to ongoing input. The longitudinal design provides an opportunity to track these dynamics across both early and extended timescales within the same subjects.

The functional consequences of the large-scale reorganization described here remain to be defined^17,62^. Although evidence indicates adult plasticity can enhance processing in spared modalities^21^, direct behavioral benefits have not been established and will require causal manipulation of the reorganized circuits. Furthermore, the extent of crossmodal recruitment of the auditory cortex predicts poorer outcomes following cochlear implantation^13,63–65^. Given the prevalence of age-related hearing loss, understanding the principles governing adult crossmodal plasticity will be essential for optimizing interventions and predicting therapeutic efficacy.

## Acknowledgements

This work was supported by the Institute for Basic Science in Korea (IBS-R015-D1); the Korea Brain Research Institute basic research program (26-BR-01-01 and 26-BR-07-01 to G.K. and 26-BR-02-02 to W.B.J.); the National Research Foundation of Korea (NRF-2021R1F1A1049434) to G.K.; and the Korea Health Industry Development Institute (KHIDI) and the Ministry of Health & Welfare of Korea (HI19C1234) to H.-J.S. We thank Dr. Hey-Kyoung Lee for critical comments on earlier versions of the manuscript, Drs. Young Rae Ji and Jisoo Han for cochlear histology and imaging, and Chanhee Lee, Junglim Lee, and Yoonsun Yang for technical assistance.

## Author contributions

H.-J.S., G.K., and S.-G.K. conceived and designed the study. H.-J.S. performed the experiments and drafted the manuscript. G.I. contributed additional fMRI data acquisition. H.-J.S. and W.B.J. analyzed the data. S.L. contributed to data organization and presentation. G.K. and S.-G.K. supervised the research. H.-J.S., W.B.J., G.K., and S.-G.K. interpreted the data and wrote the manuscript.

## Methods

### Animals

All experimental procedures were approved by the Institutional Animal Care and Use Committee (IACUC) of Sungkyunkwan University and conducted in accordance with National Institutes of Health guidelines. A total of 39 mice (7-9 weeks old; 21 males, 18 females) were used. Sensory-evoked and resting-state fMRI experiments were performed in Pou4f3^DTR/+^ mice (JAX #028673; n = 12) and wild-type littermates (n = 13). Optical intrinsic signal (OIS) imaging was performed in C57BL/6N mice (n = 5). Optogenetic fMRI experiments were performed in VGAT-ChR2-EYPF mice (JAX #014548; n = 9), which express channelrhodopsin-2 in GABAergic neurons^66^. Mice were housed in individually ventilated cages under controlled temperature and humidity on a 12 h light/dark cycle, with ad libitum access to food and water.

### Anesthesia

For all surgical procedures, fMRI, and OIS imaging, anesthesia was induced with ketamine (100 mg/kg, intraperitoneal) and xylazine (10 mg/kg, intraperitoneal) and maintained with supplemental doses (25/1.25 mg/kg) administered approximately every 40 min based on physiological monitoring^38^. Animals breathed spontaneously through a nose cone supplying a mixture of oxygen and air (1:4 ratio). Gas flow rates were 1 L/min during surgery and fMRI and 0.8 L/min during OIS imaging^44^. Oxygen saturation was maintained above 90%, and body temperature was maintained at 37 ± 0.5 °C using a feedback-controlled heating system.

### Deafening and validation

To induce deafness, diphtheria toxin (25 ng/g, intramuscular) was administered to Pou4f3^DTR/+^ mice immediately after pre-deafening MR imaging while animals remained under anesthesia. Deafness was verified using acoustic startle reflex (ASR) testing conducted before and 5-6 days after toxin injection. Broadband noise (4-16 kHz, ∼105 dB SPL, 200 ms duration) was used as the startle stimulus. For histological validation of hair cell loss, mice were transcardially perfused with saline followed by 4% paraformaldehyde. Cochleae were dissected, post-fixed overnight at 4 °C, and decalcified in 125 mM EDTA for 3-5 days. Cochlear whole-mounts were immunostained for hair cells (Myo7a), presynaptic ribbons (CtBP2), and auditory afferent fibers (Tuj1), with phalloidin used to label stereocilia, and imaged using a Leica TCS SP8 confocal microscope.

### Surgical procedures

#### Skull thinning for OIS imaging

For OIS imaging, the skull over the left cerebral cortex was thinned under anesthesia. Lidocaine (1 mg/kg, subcutaneous) was administered before scalp incision. The skull was carefully thinned using a dental drill until pial vessels were clearly visible, while the surface was kept moist with saline throughout the procedure. A thin layer of cyanoacrylate adhesive was then applied to stabilize the preparation and limit regrowth. Meloxicam (5 mg/kg, subcutaneous) was administered postoperatively. OIS imaging was performed after at least 1 week of recovery.

#### Optical cannula implantation

For optogenetic experiments, optic cannulas (105 µm, 0.22 NA) were implanted bilaterally in primary auditory cortex (AP −2.6 mm, ML ±4.3 mm, depth 0.7 mm) under anesthesia, as previously described^67^. Briefly, small burr holes were made at the target coordinates, and cannulas were lowered into A1 and secured using silicone sealant and dental cement. Experiments were performed after at least 10 days of recovery.

### Sensory stimulation and optogenetic silencing

During sensory-evoked fMRI, somatosensory and visual stimuli were delivered in alternating trials. For OIS imaging and optogenetic fMRI, only forepaw stimulation was used. Somatosensory stimulation was delivered through a pair of needle electrodes inserted into the left forepaw (4 Hz, 0.5 ms pulse width, 0.7 mA). Visual stimulation consisted of white LED pulses delivered through a branching fiber-optic patch cord positioned approximately 1 cm in front of both eyes (5 Hz, 10 ms pulse width, ∼15 lux).

Optogenetic silencing was achieved by delivering blue light (473 nm) through implanted fibers (20 Hz, 10 ms pulse width), as previously described^68^. During sensory-evoked fMRI with optogenetic silencing, laser stimulation was applied continuously throughout each run, maintaining A1 in either a silenced (laser-on) or control (laser-off) state. Laser output was calibrated to ∼3 mW at the fiber tip, corresponding to a time-averaged power density of 69.3 mW/mm². To minimize MR susceptibility artifacts, the skull was sealed and light leakage was prevented by shielding the fiber connection. All sensory stimuli were triggered by a pulse generator and synchronized with MRI acquisition.

### MRI data acquisition

All MRI experiments were performed on a horizontal-bore 15.2 T MRI scanner (Bruker BioSpec) equipped with an actively shielded 6-cm gradient (maximum strength, 100 G/cm, rise time, 110 µs). A custom-built 15 mm inner diameter surface coil was positioned over the mouse head to cover the cortex. Magnetic field homogeneity was optimized over an ellipsoidal volume covering the entire brain using FASTMAP shimming.

Anatomical images were acquired using a FLASH sequence (TR/TE = 384.06/3.34 ms, flip angle = 30°, field of view = 15.8 × 7.65 mm², matrix size = 256 × 128, 4 averages, spatial resolution = 62 × 60 × 500 µm³, 20 contiguous coronal slices). These images were used for spatial normalization to atlas space. BOLD fMRI data were acquired using a single-shot gradient-echo echo-planar imaging (EPI) sequence (sampling frequency = 300 kHz, TR/TE = 1000/11.5 ms, flip angle = 50°, field of view = 15.8 × 7.65 mm², matrix = 120 × 58, spatial resolution = 132 × 132 × 500 µm³, 20 contiguous coronal slices). For sensory-evoked fMRI, each trial consisted of 200 volumes acquired using a two-block design comprising 40 s baseline, 20 s stimulation, 60 s rest, followed by a second 20 s stimulation and 60 s rest period. The inter-trial interval was approximately 1 min, and each session included 10 trials per stimulus condition (forepaw and visual). Resting-state fMRI was acquired using the same imaging parameters in two runs of 300 volumes (5 min each). Sensory-evoked and resting-state data were acquired from the same animals before and 1 week after DT administration.

### fMRI preprocessing and analysis

MRI data were analyzed using Analysis of Functional Neuroimages package (AFNI)^69^, FMRIB Software Library (FSL)^70^, Advanced Normalization Tools (ANTs)^71^ and custom MATLAB code (MathWorks, Natick, USA).

#### Sensory-evoked fMRI

Sensory-evoked fMRI data were preprocessed using a pipeline described previously in detail^39^. Preprocessing included slice-timing correction, motion realignment to correct for intra- and inter-scan head motion, linear detrending, and normalization of voxel time courses to the mean baseline signal. Trials belonging to the same stimulus condition were averaged to improve signal-to-noise ratio. Functional images were coregistered to anatomical images and spatially normalized to the Allen Mouse Brain Atlas using deformation fields derived from anatomy-to-atlas registration. Normalized functional images were spatially smoothed with a Gaussian kernel (FWHM = 0.2 mm).

Activation maps for individual animals were generated using a general linear model (GLM), in which stimulus paradigms were convolved with a ketamine/xylazine-specific hemodynamic response function. Atlas-based regions of interest (ROIs) were defined across somatosensory (S1FL, S2, M1, VP), visual (V1, V2, LGd/LP, SC), auditory (A1, MG, IC), and association (ACC, RSC, PPC) areas, with the reticular thalamic nucleus (RT) used as a control region. For somatosensory stimulation, ROIs were defined in the hemisphere contralateral to the stimulated forepaw, whereas bilateral ROIs were used for visual stimulation. For optogenetic fMRI, a subset of ROIs (S1FL, S2, M1, VP, V1, and A1) was analyzed. Time courses were extracted from each ROI, and stimulus-evoked responses were quantified as percent signal change averaged over the stimulation period.

#### Resting-state fMRI

Resting-state data were analyzed using a pipeline adapted from previous work^72^. Preprocessing steps included slice-timing correction, motion realignment, linear detrending, voxel-wise normalization, and spatial smoothing with a Gaussian kernel (FWHM = 0.2 mm). To improve the reliability of functional connectivity estimates, resting-state fMRI data were additionally despiked, regressed for 12 motion-related confounds (six motion parameters and their temporal derivatives), and band-pass filtered (0.01–0.2 Hz). Group-level independent component analysis (ICA) was performed using MELODIC in FSL^73^ after temporal concatenation across animals. The dimensionality was set to 20 components, and a threshold of 0.5 was used to define significant components. From the ICA-derived spatial maps, ten components were selected for further analysis as anatomically plausible large-scale resting-state networks based on their spatial correspondence to the Allen Mouse Brain Atlas. These included somatosensory (SS#1: S1_N_ + S2; SS#2: bilateral S1_BF_ + S1_FL_), motor (Mot: M1 + M2 + S1_FL_), visual (Vis: V1 + V2), auditory and temporal association (Aud: AUD + TEa), default-mode-like (DM: ACC + RSC), hippocampal (HC#1: HIP; HC#2: RHP), dorsal striatum (Str: STRd), and olfactory/lateral septal (Olf: PIR + LS) networks. Subject-specific component time courses were estimated using dual regression. Functional connectivity matrices were then constructed by calculating Pearson correlation coefficients between all pairs of components for each animal and condition. Correlation coefficients were converted using Fisher’s r-to-z transformation before statistical comparison.

### CBV-weighted Optical intrinsic signal (OIS) imaging

Cerebral blood volume (CBV)-weighted OIS imaging was performed in naïve mice under alternating noise-on and noise-off conditions to assess whether EPI scanner noise affects somatosensory-evoked hemodynamic responses. In the fMRI protocol, EPI-related scanner noise reached ∼115 dB with a peak power spectral density near 1.3 kHz. To reproduce this acoustic environment, recorded scanner noise was played back during OIS imaging. Because OIS, like BOLD fMRI, reflects stimulus-evoked hemodynamic responses, it was used as an independent test of this potential confound^74^.

OIS data were acquired through the thinned-skull preparation using an MVX-10 microscope (Olympus) equipped with a 580 nm band-pass filter and illumination from a 572 nm LED (isosbestic point of hemoglobin) to measure total hemoglobin/CBV changes^74^. Images were collected at 2 frames/s with a resolution of 540 × 372 pixels over an 8.6 × 5.9 mm² field of view and downsampled to 270 × 186 pixels by 2 × 2 spatial binning. Each trial consisted of 20 s forepaw stimulation followed by 60 s rest, and 12 trials were acquired per animal for each noise condition.

For analysis, normalized activation maps were calculated as ΔR(t)/R, where ΔR(t) = R(t) – R_0_ and R_0_ was the baseline reflectance averaged from −15 to −5 s before stimulus onset. A 1 × 1 mm² square ROI was centered on the activation focus defined from the mean activation map during the 10–20 s window and applied consistently across conditions. Time courses were extracted by averaging pixel intensities within the ROI. Because increases in CBV decrease reflectance, the sign of ΔR(t)/R was inverted for visualization and quantification. Mean CBV responses were quantified over the 10–20 s post-stimulus window.

### Statistical analysis

Data are presented as mean ± standard error of the mean (SEM), unless otherwise stated. Within-subject comparisons, including pre- versus post-DT measurements, noise-off versus noise-on OIS responses, and sensory-evoked fMRI responses with and without optogenetic silencing, were performed using paired two-tailed Student’s t-tests. A P value < 0.05 was considered statistically significant.

---

**Supplementary Figure 1.**
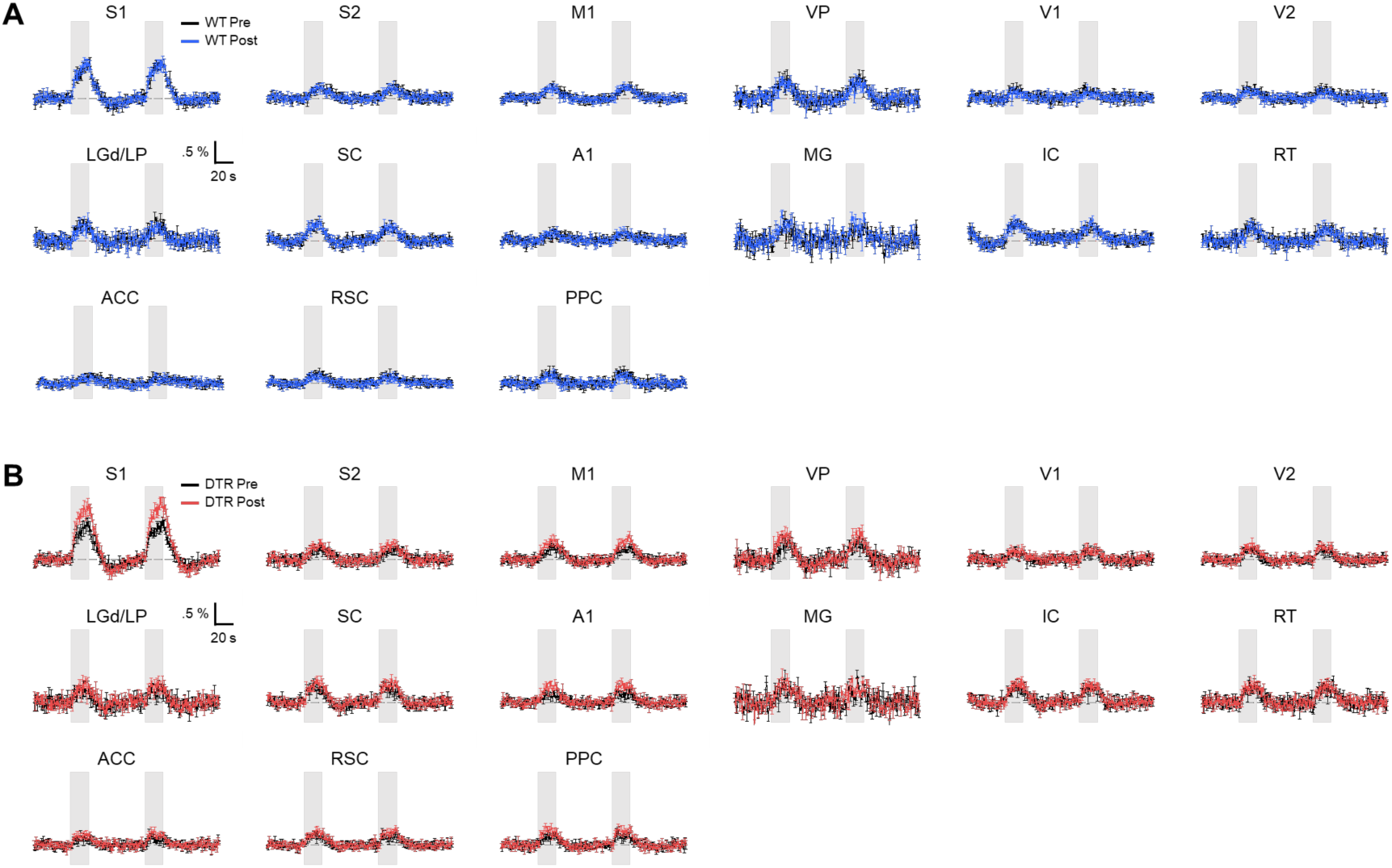
Time-course analysis of somatosensory-evoked BOLD responses across all 15 ROIs in WT (n = 13 mice) and DTR mice (n = 12 mice). Temporal response profiles were preserved in both groups, while DTR mice exhibited increased response amplitudes after deafening.

**Supplementary Figure 2.**
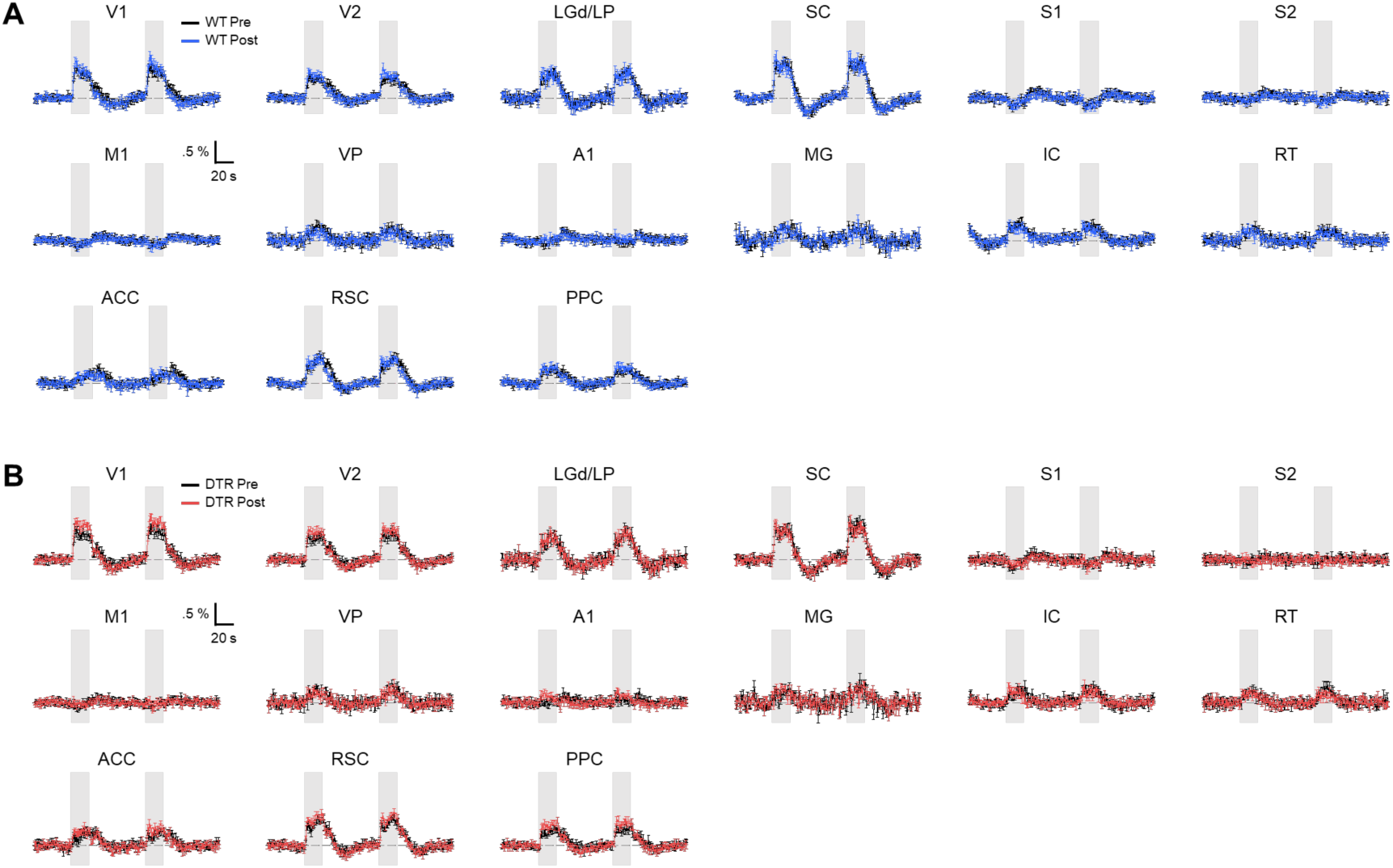
Time-course analysis of visual-evoked BOLD responses across all ROIs in WT (n = 13 mice) and DTR (n = 12 mice) mice. Visual stimulation evokes preserved temporal dynamics with increased response amplitudes in cortical regions after deafening, without shifts in response latency or duration.

