## Supplemental Tables 1-6 for "Deafness rapidly reorganizes functional brain networks in adult mice"

**Supplementary Table 1**
Somatosensory-evoked BOLD responses in wild-type mice

| ROI | Pre (%) | Post (%) | p-value | Direction |
|---|---|---|---|---|
| S1FL | 0.51 ± 0.05 | 0.57 ± 0.06 | > 0.05 | - |
| S2 | 0.12 ± 0.02 | 0.16 ± 0.02 | > 0.05 | - |
| M1 | 0.14 ± 0.01 | 0.18 ± 0.02 | > 0.05 | - |
| VP | 0.30 ± 0.03 | 0.24 ± 0.04 | > 0.05 | - |
| V1 | 0.13 ± 0.02 | 0.09 ± 0.02 | > 0.05 | - |
| V2 | 0.14 ± 0.02 | 0.11 ± 0.01 | > 0.05 | - |
| LGd/LP | 0.25 ± 0.03 | 0.23 ± 0.04 | > 0.05 | - |
| SC | 0.23 ± 0.02 | 0.28 ± 0.03 | > 0.05 | - |
| A1 | 0.10 ± 0.02 | 0.12 ± 0.02 | > 0.05 | - |
| MG | 0.17 ± 0.04 | 0.26 ± 0.03 | > 0.05 | - |
| IC | 0.26 ± 0.02 | 0.27 ± 0.02 | > 0.05 | - |
| ACC | 0.09 ± 0.02 | 0.09 ± 0.01 | > 0.05 | - |
| RSC | 0.14 ± 0.02 | 0.13 ± 0.02 | > 0.05 | - |
| PPC | 0.17 ± 0.02 | 0.13 ± 0.02 | > 0.05 | - |
| RT | 0.21 ± 0.03 | 0.16 ± 0.02 | > 0.05 | - |

Values are shown as mean ± SEM.
No significant changes were observed in wild-type mice (all $p > 0.05$).

**Supplementary Table 2**
Somatosensory-evoked BOLD responses in deafened (Pou4f3$^{DTR/+}$) mice

| ROI | Pre (%) | Post (%) | p-value | Direction |
|---|---|---|---|---|
| **S1FL** | 0.56 ± 0.07 | **0.86 ± 0.07** | *** | ↑ |
| **S2** | 0.15 ± 0.03 | **0.24 ± 0.02** | * | ↑ |
| **M1** | 0.17 ± 0.02 | **0.31 ± 0.03** | ** | ↑ |
| **VP** | 0.25 ± 0.03 | **0.38 ± 0.02** | ** | ↑ |
| V1 | 0.13 ± 0.02 | 0.16 ± 0.02 | > 0.05 | - |
| V2 | 0.17 ± 0.02 | 0.22 ± 0.02 | > 0.05 | - |
| LGd/LP | 0.21 ± 0.04 | 0.28 ± 0.03 | > 0.05 | - |
| SC | 0.26 ± 0.04 | 0.33 ± 0.04 | > 0.05 | - |
| **A1** | 0.14 ± 0.04 | **0.28 ± 0.03** | ** | ↑ |
| MG | 0.20 ± 0.06 | 0.27 ± 0.03 | > 0.05 | - |
| IC | 0.25 ± 0.03 | 0.30 ± 0.04 | > 0.05 | - |
| **ACC** | 0.11 ± 0.02 | **0.19 ± 0.01** | ** | ↑ |
| **RSC** | 0.16 ± 0.01 | **0.23 ± 0.02** | ** | ↑ |
| **PPC** | 0.16 ± 0.02 | **0.27 ± 0.03** | ** | ↑ |
| RT | 0.22 ± 0.03 | 0.26 ± 0.02 | > 0.05 | - |

Values are shown as mean ± SEM.
* p < 0.05; **: p < 0.01; ***: p < 0.001 (paired comparison).

**Supplementary Table 3**
Visual-evoked BOLD responses in wild-type mice

| ROI | Pre (%) | Post (%) | p-value | Direction |
|---|---|---|---|---|
| S1FL | -0.13 ± 0.03 | -0.11 ± 0.02 | > 0.05 | - |
| S2 | -0.04 ± 0.03 | -0.04 ± 0.02 | > 0.05 | - |
| M1 | -0.08 ± 0.03 | -0.07 ± 0.01 | > 0.05 | - |
| VP | 0.20 ± 0.03 | 0.16 ± 0.02 | > 0.05 | - |
| V1 | 0.51 ± 0.04 | 0.56 ± 0.07 | > 0.05 | - |
| V2 | 0.37 ± 0.03 | 0.43 ± 0.04 | > 0.05 | - |
| LGd/LP | 0.47 ± 0.04 | 0.43 ± 0.07 | > 0.05 | - |
| SC | 0.66 ± 0.07 | 0.65 ± 0.09 | > 0.05 | - |
| A1 | -0.03 ± 0.03 | 0.02 ± 0.02 | > 0.05 | - |
| MG | 0.19 ± 0.03 | 0.17 ± 0.04 | > 0.05 | - |
| IC | 0.28 ± 0.05 | 0.24 ± 0.04 | > 0.05 | - |
| ACC | 0.13 ± 0.03 | 0.16 ± 0.03 | > 0.05 | - |
| RSC | 0.41 ± 0.05 | 0.42 ± 0.04 | > 0.05 | - |
| PPC | 0.23 ± 0.03 | 0.30 ± 0.03 | > 0.05 | - |
| RT | 0.22 ± 0.03 | 0.26 ± 0.02 | > 0.05 | - |

Values are shown as mean ± SEM.
No significant changes were observed in wild-type mice (all p > 0.05).

**Supplementary Table 4**
Visual-evoked BOLD responses in deafened ($Pou4f3^{DTR/+}$) mice

| ROI | Pre (%) | Post (%) | p-value | Direction |
|---|---|---|---|---|
| S1FL | -0.08 ± 0.03 | -0.08 ± 0.02 | > 0.05 | - |
| S2 | -0.01 ± 0.03 | -0.02 ± 0.02 | > 0.05 | - |
| M1 | -0.04 ± 0.03 | -0.03 ± 0.02 | > 0.05 | - |
| VP | 0.18 ± 0.03 | 0.18 ± 0.02 | > 0.05 | - |
| **V1** | 0.51 ± 0.04 | **0.69 ± 0.04** | ** | ↑ |
| **V2** | 0.44 ± 0.04 | **0.55 ± 0.03** | * | ↑ |
| LGd/LP | 0.42 ± 0.06 | 0.43 ± 0.05 | > 0.05 | - |
| SC | 0.62 ± 0.10 | 0.59 ± 0.07 | > 0.05 | - |
| **A1** | 0.03 ± 0.03 | **0.13 ± 0.03** | * | ↑ |
| MG | 0.20 ± 0.04 | 0.21 ± 0.04 | > 0.05 | - |
| IC | 0.22 ± 0.05 | 0.21 ± 0.05 | > 0.05 | - |
| **ACC** | 0.18 ± 0.03 | **0.30 ± 0.03** | * | ↑ |
| **RSC** | 0.44 ± 0.04 | **0.53 ± 0.03** | * | ↑ |
| **PPC** | 0.28 ± 0.03 | **0.41 ± 0.04** | ** | ↑ |
| RT | 0.19 ± 0.03 | 0.15 ± 0.02 | > 0.05 | - |

Values are shown as mean ± SEM.
* p < 0.05; **: p < 0.01 (paired comparison).

**Supplementary Table 5**
Somatosensory-evoked BOLD responses during acute auditory cortex suppression

| ROI | W/O A1s | W/ A1s | p-value |
|---|---|---|---|
| S1FL | 0.68 ± 0.10 | 0.64 ± 0.09 | > 0.05 |
| S2 | 0.18 ± 0.04 | 0.21 ± 0.03 | > 0.05 |
| M1 | 0.27 ± 0.05 | 0.22 ± 0.05 | > 0.05 |
| VP | 0.33 ± 0.06 | 0.31 ± 0.06 | > 0.05 |
| V1 | 0.14 ± 0.09 | 0.13 ± 0.05 | > 0.05 |
| A1 | 0.12 ± 0.03 | 0.14 ± 0.03 | > 0.05 |

Values are shown as mean ± SEM.
No significant changes were observed in VGAT-ChR2 mice (all $p > 0.05$).

**Supplementary Table 6**
Resting-state functional connectivity changes after deafening

| Network pair | Pre (z) | Post (z) | Δ(Post-Pre) | p-value |
|---|---|---|---|---|
| SS#1 to Vis | -0.03 ± 0.03 | 0.04 ± 0.04 | 0.07 ± 0.03 | * |
| Mot to Vis | 0.08 ± 0.04 | 0.17 ± 0.05 | 0.09 ± 0.03 | * |
| Aud to Vis | 0.18 ± 0.05 | 0.26 ± 0.04 | 0.08 ± 0.03 | * |
| DM to SS#2 | 0.19 ± 0.04 | 0.39 ± 0.07 | 0.20 ± 0.08 | * |
| DM to Vis | 0.29 ± 0.05 | 0.49 ± 0.06 | 0.20 ± 0.07 | * |
| DM to Aud | 0.11 ± 0.03 | 0.20 ± 0.03 | 0.10 ± 0.03 | ** |

Values are shown as mean ± SEM.
Δ(Post-Pre) values were computed per animal.
* $p < 0.05$; **: $p < 0.01$ (paired comparison).
No significant changes were observed in wild-type mice for any network pair (all $p > 0.05$).
